# Evaluation of Antinociceptive and Anti-Inflammatory Activities of Solvent Fraction of the Roots of *Echinops kebericho* Mesfin (Asteraceae) In Mice Model

**DOI:** 10.1101/2023.03.06.531449

**Authors:** Tesfaye Yimer Tadesse, Samuel Berihun Dagnew, Tesfagegn Gobezie Yiblet, Getu Tesfaw Addis, Zemene Demelash Kiflie

## Abstract

**Background:** Since ancient times, pain and inflammation have been treated using herbal remedies, which are essentially a stockroom of phytochemical components. Due to the numerous adverse effects of the already available anti-pain and anti-inflammatory medications, the search for new potential pharmaceuticals used to relieve pain and inflammation from natural sources is an ongoing process. The present study was therefore, aimed at investigating the antinociceptive and antiinflammatory activities of the solvent fractions of the roots of *E. kebericho* M. in mice model.

**Methods:** Successive maceration was used as a method of extraction using solvents of increasing polarity: methanol and water. The crude extract was then further fractionated using distilled water, ethyl acetate, and chloroform. Each solvent fraction was then evaluated for its peripheral analgesic activities using an acetic acid-induced writing test and central analgesic activities using the hot plate method. The acute and chronic anti-inflammatory activities of the solvent fractions were detected using carrageenan induced paw edema and cotton pellet ear granuloma respectively. The detected doses were 100mg/kg, 200mg/kg, and 400mg/kg. The positive control groups received ASA (150mg/kg) for the writing test, morphine (10mg/kg) for the hot plate method, diclofenac Na for carrageenan induced paw edema and dexamethasone (10mg/kg) for granuloma, while the negative control group received distilled water.

**Result:** EA fraction at all test doses employed (100mg/kg, 200mg/kg and 400mg/kg) showed statistical significant (p < 0.05, p < 0.01, p < 0.001 respectively) analgesic effects in both chemical and thermal induced pain stimuli in dose dependant manner. Likewise, EA fraction also exhibited anti-inflammatory activities on carrageenan induced paw edema and cotton pellet-induced granuloma in a dose-dependent manner. The AQ fraction on the other hand produced statistical significant (p < 0.05, p < 0.012) analgesic and anti-inflammatory activities at the doses of 200mg/kg and 400mg/kg, while the CH fraction exhibited statistical significant (p < 0.05) analgesic and anti-inflammatory activity at the dose of 400mg/kg.

**Conclusion:** In general, the data obtained from the present study elucidated that the solvent fractions possessed significant analgesic and anti-inflammatory activities and recommended further investigations.

## INTRODUCTION

Pain and inflammation are the most common manifestations of variety of conditions that impact many individuals globally, these clinical conditions are the two most important areas of global scientific study (1). Whenever there is actual or potential tissue damage, pain is always a subjective, unpleasant sensory and emotional experience. It is often triggered by painful stimuli, which are subsequently transmitted to the central nervous system (CNS) and recognized as such (1, 2). It is an immediate reaction to an undesirable occurrence linked to tissue damage, like an injury, inflammation, cancer, osteo, and rheumatoid arthritis. In addition, severe pain might arise suddenly (trigeminal neuralgia), persist for a long time after the initial injury has healed (phantom limb pain), or be brought on by a harmless stimulus (allodynia) (1, 3, and 4). It serves as the body’s means of defense against harm (5). Similar to how it conducts body rescue functions, inflammation also neutralizes, destroys, and dilutes dangerous substances. Even though, the inflammatory processes, that are the body’s defense mechanism, aid in the removal of pathogens and other harmful stimuli and start the healing process for the inflammatory reaction, untreated and long term inflammatory processes harm the body organs and resulted organ failures and even death (6, 7). Both pain and inflammations are the most common reasons for patients seeking medical interventions (8).

Despite the availability of medically proven ant pain and anti-inflammatory agents, pain and inflammation continues to rank among the most challenging health issues, impacting more than 80% of adults worldwide. They are frequently viewed as a significant clinical, social, and economic problem in communities all over the world. Untreated chronic pain, which harms both physical and psychological health, is the most prevalent issue. Many people seek medical assistance to manage their pain because it is the most prevalent and common cause of mental impairments, accounting for approximately 35 million office visits annually (8, 9). Pain is the most serious public health concern because it affects all populations, regardless of age, gender, income, race/ethnicity, or geography. If not properly controlled and managed, pain can have many serious health consequences, including depression, the inability to work, strained social relationships, and suicidal thoughts (10).

Inflammation which is mediated by both the innate and adaptive immune systems is a physiological response of living tissues to injury (11). Although the inflammatory response is essential for host defense, it is very much a double-edged sword which can lead to an organ failure and/or death unless it is properly controlled and managed (12, 13).

Currently, different classes of drugs are available for the pharmacological management of pain and inflammation. However, their effectiveness against pain and different inflammatory conditions is limited due to different problems including unaffordability, limited accessibility, adverse drug reactions, and many medicines that are not effective as expected in all patients (14, 15). Tolerance and dependence is another challenge, particularly with the chronic use of opioids. Prolonged uses of NSAIDs also lead to different untoward effects such as gastric irritation, gastric ulcer, alterations in renal function, effects on blood pressure, hepatic injury, and platelet inhibition which may result in increased bleeding (14, 15, 16). As a result, more research into medicinal plants that have been reported to be useful in the treatment of pain and inflammation is required.

Due to their easy accessibility, availability, low cost, and safety, herbal medicines are used to replace or assist conventional therapies in the treatment of various diseases all over the world so far. Thus, it seems to be worthwhile searching for traditional medicinal plants which possess potential anti-inflammatory and antinociceptive activities that are used to heal different inflammatory and pain-related disorders by traditional healers (17, 18). It is known that medicinal plants have a large diversity of secondary metabolites with different biological activities which justifies the research on the pharmacological properties of plant species and their potential uses in drug development (17, 19, 20). Conventionally many plant species are used as analgesic and antiinflammatory agents in Ethiopia folk medicine. Some of these plants include *Echinopskebericho* (54), *Ocimumlamiifolium, Deciliterlaxata, CrotonMacrostachys, Vernoniaamygdalina, Caricapapaya, Eucalyptusglobules, AlliumSativam, Echinopsmaccrochaetus, Schinusmolle*, and *Withaniasomnifera* (55).

As stated (21) *E. kebericho* M. has been used for the relief of different infectious and non-infectious diseases in different preparations. The analgesic and anti-inflammatory effects of 80% methanol root crude extract of the plant possessed significant analgesic activities in both chemical-induced peripheral pain and thermal-induced central pain stimuli. Besides, the anti-inflammatory potential of the crude extract was detected in acute and sub-acute inflammatory phases and it possessed statistically significant ant-inflammatory potential (21).

Even though the plant *E. kebericho* M. is frequently reported for its antinociceptive and antiinflammatory potentials by traditional healers in different parts of Ethiopia, and the previous reports showed that the crude extract exhibited statistical significant analgesic and antiinflammatory activities, further investigation through the solvent fraction is deemed prudent to illustrate the most active plant constituents for its anti-inflammatory and analgesic activities. The aim of the present study was, therefore, to investigate the analgesic and anti-inflammatory activities of solvent fractions of the roots of *E. kebericho* M. in a mice model.

## MATERIALS AND METHODS

### Materials and Instruments

Rotary evaporator (yamato, Japan), lyophilizer (OPERON, OPR-FDU-5012, Korea), digital plethysmometer (Ugo Basile-Cat no 7140, Italy) electronic balance (KERN-ALJ 220–4, Germany), tissue Drying Oven (Medite – Medizin technik, Germany), syringes with needles, feeding tube, hot Plate (Orchid Scientific, India.) were used with their respective function.

### Drugs and chemicals

Carrageenan (Sigma Chemicals Co., St Louis, USA), formalin (Taflen Industry, Ethiopia), normal saline (H. R. Leuven, Belgium), distilled water (Ethiopian Pharmaceutical Manufacturing Factory, Ethiopia), absolute methanol (Indenta chemicals, India) and glacial acetic acid (Sigma-Aldrich laborchemikalien, Germany), indomethacin (Cadila pharmaceuticals Ethiopia), aspirin, diclofenac, and morphine (Ethiopian Pharmaceutical Manufacturing Factory, Ethiopia) obtained from the respective vendors were used in the experiment.

### Collection, Identification, and Preparation of Plant Materials

The roots of *E. kebericho* were collected from around Debre tabor town South Gondar Zone of the Amhara regional state which is located 667km far from North West Addis Ababa, the capital city of Ethiopia. Then the plant was authenticated by Botanists in the Department of Biology, College of Natural and Computational Sciences, Debre Tabor University where a specimen with voucher number 001TYT/2021 was deposited for future reference.

### Extract Preparation

After collection, the roots of the plant will be washed with tape water to remove dust and any other debris present on them. Then the roots of E. *kebericho* was air dried under a shaded area at room temperature and subjected to be pulverized using a mortar and pestle to get a coarse powder used for the extraction. After all, a total amount of 5kg powdered root was macerated using 99.9% Absolute methanol with distilled water in the ratio of 4:1 for about 72 hrs. The maceration was undertaken with occasional shaking using mini orbital shaker being tuned to 120 rpm for 72 hrs. at room temperature. Then, the extract was filtered first using muslin cloth and then using Whatman filter paper No 1. Filtration and collection of the extract were repeated three times with the whole extraction taking a total of 9 days. After the extraction was completed, methanol and water were evaporated under vacuum using rotary vapor and oven at 40°C. The resulting solution was placed in a deep freezer operating at -20°C till it formed solid ice and then the remaining solvent (water) will be removed using lyophilizer. After crude extraction was completed, the resulting extract was further subjected to be fractionated using solvents of different polarity i.e. ethyl acetate, distilled water, and chloroform. After all, the resulting fractions were within a deep refrigerator (4°c) till the commencement of the main procedure (3, 44).

### Experimental Animals

Healthy adult Swiss Albino mice of either sex (25-35g body weight and 6–8 weeks of age) were purchased from Ethiopian Health and Nutrition Research Institute (EHNRI). The animals were kept in cages at room temperature on a 12 hrs. light / dark cycle with access to standard laboratory pellets and water *ad libitum*. All the experimental animals were allowed to be acclimatized to the laboratory condition for a week before the commencement of the experiment. All animals used in this study were handled in accordance with the internationally accepted standard guidelines for use of laboratory animals

### Preliminary Phytochemical Screening

The analgesic and anti-inflammatory activities of the solvent fraction of the roots of *E. kebericho* were investigated in relation to the presence or lack of secondary metabolites using a conventional phytochemical screening test. Thus, the test for alkaloids, saponins, flavonoids, phenols, steroid, anthraquinone, glycosides, and tannins were performed using standard test procedures (44, 54).

### Animal Grouping and Dosing

Swiss albino mice of either sex (25-35g BW) were randomly divided into eleven groups of six mice per group. Group, I was assigned as negative control and received vehicles. Group II served as positive control and was treated with standard drugs; morphine (10mg/kg, s.c) for hot plate test, indomethacin (25 mg/kg, p.o) cotton pellet granuloma, ASA (150 mg/kg, p.o) for writhing. Groups III-XI was used as test groups and was given the solvent fractions at the dose of 100mg/kg, 200mg/kg, and 400mg/kg respectively. Doses were selected based on an acute toxicity study done previously (56, 65).

### Antinociceptive Activity Test

#### Acetic Acid Induced Writhing Method

This method was conducted to detect the peripheral antinociceptive activities of the solvent fractions of the roots of *E. kebericho* in mice and was performed by randomly dividing overnight fasted mice of either sex (25-35g) with free water access into twelve groups, consisting of six mice per group. Groups I -IX were assigned as test groups and were given different doses of *E*.*kebericho* solvent fraction (100mg/kg, 200mg/kg, and 400mg/kg) which was determined under acute toxicity and dosing done in the previous studies (65, 66). Group X was used as negative control and was given a vehicle (distilled water). The eleventh (positive control) group was given 150mg/kg dose of the standard drug ASA. 0.6% v/v of acetic acid (10ml/kg, i.p) was injected into all groups of mice an hour just after the mice were given the extract, vehicle, and the standard drug with their respective doses. Analgesic activity of each fraction was assessed by counting the numbers of writhing which consists of contraction of the abdominal muscle together with stretching of the hind limbs for 30 min after a latency period of 5 min. A reduction in the number of writhes as compared to the control group was considered as evidence for the analgesic potential of each fraction, and was expressed as percent inhibition of writhing, according to the following formula (71).

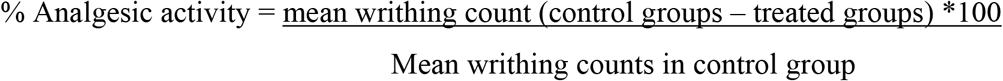

### Hot Plate Method

This method was carried out to evaluate the central antinociceptive potentials of the roots *E. kebericho* solvent fraction and was performed by introducing the mouse into an open-ended cylindrical space with a floor consisting of a metallic plate that was heated by a thermode. A plate was heated to a constant temperature of 55°C ± 1°C producing the behavioral components that were measured in terms of their reaction times, namely paw licking, withdrawal of the paw, and jumping. All responses were considered to be supraspinally integrated responses. Each mouse was placed on a hot plate individually with a cut-off time of 15s to avoid lesions to the animals’ paws. The latency to lick the paw or jump from the hot plate was recorded as the reaction time. The reaction times were noted at 0 and 30, 60, 90 and 120 min after the administration of vehicle (distilled water 10ml/kg, standard drug (morphine10mg/kg) and 100mg/kg, 200mg/kg and 400mg/kg of each solvent fraction. Percentage increase in reaction time or pain threshold inhibition was calculated using the formula as follows (72, 73).

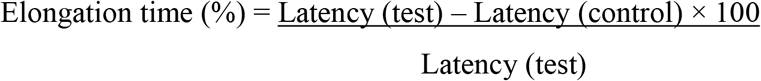

### Evaluation of Anti-inflammatory Activity of each solvent fraction

#### Carrageenan Induced Paw Edema

This procedure was carried out by giving mice that had been fasting overnight free access to water and inducing an acute inflammation in their paws. Carrageenan (1% w/v in normal saline, 0.05mL) was administered to the mice via injection into the left hind paw’s plantar side. To ensure that each mouse’s leg could be submerged to the same depth in the plethysmometer’s measuring chamber, the skin around the lateral malleolus of each animal was marked just before inflammation was induced. One hour after giving the appropriate groups of mice the solvent fractions, the vehicle, and the standard medication, carrageenan was administered. Inflammation was expressed in terms of mL i.e., displacement of water by edema using a digital plethysmometer at time 0, 1, 2, 3, 4 after carrageenan injection. The percentage inhibition of edema was calculated in terms of edema volume using the following formula (73).

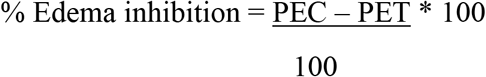

Where; PEC paw edema in control group

PET paw edema in test group

### Cotton Wool Induced Granuloma

Cotton wool granuloma which is used to detect anti-proliferative activities of natural products was applied to detect the anti-inflammatory effects of solvent fractions of the roots E. *kebericho* in chronic inflammation. The procedure was performed by provoked foreign body granuloma in mice by subcutaneous implantation of pellets of compressed cotton. After several days, histologically giant cells and undifferentiated connective tissue was observed besides the fluid infiltration. The amount of newly formed connective tissue was then measured by weighing the dried pellets after removal. The experimental mice were anesthetized with ether and the back skin was shaved and disinfected with 70% ethanol. An incision was made in the lumbar region. By blunted forceps subcutaneous tunnels will be formed and a sterilized cotton pellet will be placed on both sides in the scapular region. The pellets formed from raw cotton which produces a more pronounced inflammation than bleached cotton. The animals were treated for 7 days. Then, the animals were sacrificed; the pellets were prepared and dried until the weight remains constant. The net dry weight, i.e. after subtracting the weight of the cotton pellet was determined (74, 75).

### Statistical analysis

The data that was obtained from the experiments were expressed as mean ± SEM. The results were statistically analyzed using one-way ANOVA followed by Post Hoc Tukey-tests for multiple group comparison with SPSS version 25 software. The results were considered significantly different at p < 0.05.

## Results

### Preliminary phytochemical screening

The preliminary phytochemical screening’s goal was to investigate the different kinds of secondary metabolites present in each solvent fraction based on qualitative color changes in test reagents, which could provide information on how analgesic and anti-inflammatory effects might relate to the presence of different active phytoconstituents. The solvent fractions from the roots of *E. kebericho* showed the presence of numerous secondary metabolites based on preliminary phytochemical screening assays. The only substances found in the chloroform fraction are steroids and alkaloids. Tanning agents, flavonoids, saponins, and cardiac glycosides were present in the aqueous fraction. The richest fraction, however, was the ethyl acetate one since it contained tannins, alkaloids, saponins, terpenoids, and anthraquinones.

### Antinociceptive Activity

#### Acetic acid-induced writhing test

This method was conducted to evaluate the peripheral anti-nociceptive effects of the solvent fractions of the roots of *E*.*kebericho* to chemical-induced pain stimuli. In this method, the number of writhes (in 15 min) was highest in the negative controlled group (distilled water treated mice) (64.17 ± 0.40) and lowest in AF 400 treated mice (23.2 ± 0.95) (**Table 1**). The mean writhing reduction produced by the middle (200mg/kg) and the highest (400mg/kg) doses of the ethyl acetate, aqueous, and chloroform fractions of *E. kebericho* were statistically significant as compared to the negative control (p < 0.01 and 0.001 respectively) (**Table 1)**. All the employed test doses of each solvent fraction produced increased inhibition of the numbers of writhing with maximum inhibition observed at the highest doses (400 mg/kg) (p < 0.01 for EA, p < 0.001 for AQ and CH fractions). The dose-dependent writhing reduction was observed in each solvent fraction. The extent of reduction of writhing in the different doses of each solvent fraction was different, i.e. (p < 0.001) for 400 mg/kg and (p < 0.01) for 200 mg/kg ethyl acetate fraction. The lowest doses (100mg/kg) of each solvent fraction showed no significant effect on chemical-induced writhing as compared to the negative control groups. On the other hand, the standard drug (ASA 150mg/kg) produced a high percentage of writhing reduction (40.30 %) as compared to other groups. Ethyl acetate fraction produced a 33.08 % writhing reduction while AQ and CH fractions produced a percentage writhing reduction of 29.35% and 25.37% respectively (**Table 1**)

**Table 1.**
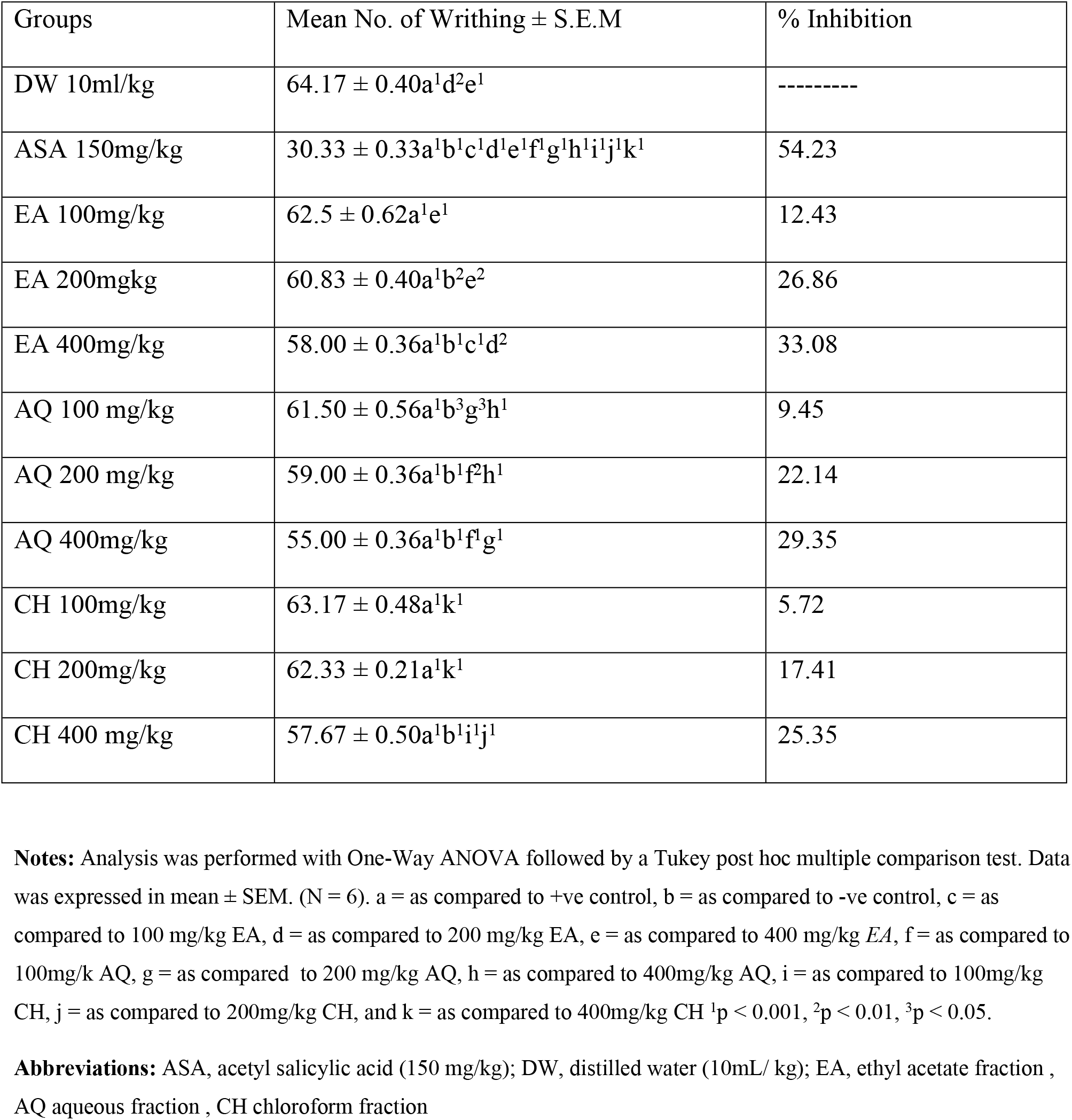
Effects of solvent fractions of the roots of *Echinops kebericho* on Acetic Acid-Induced Writhing in Mice

**Table 1.**
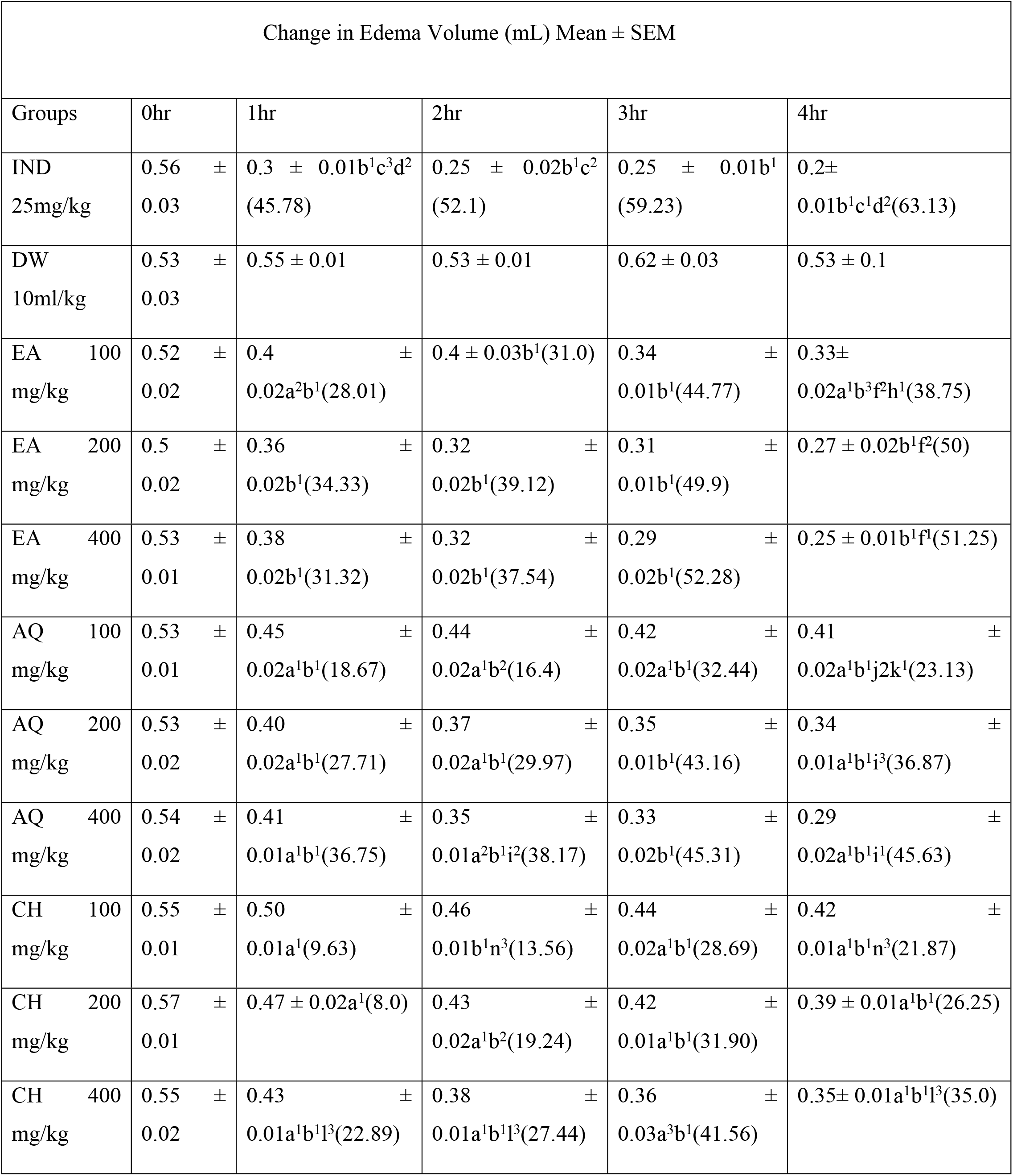
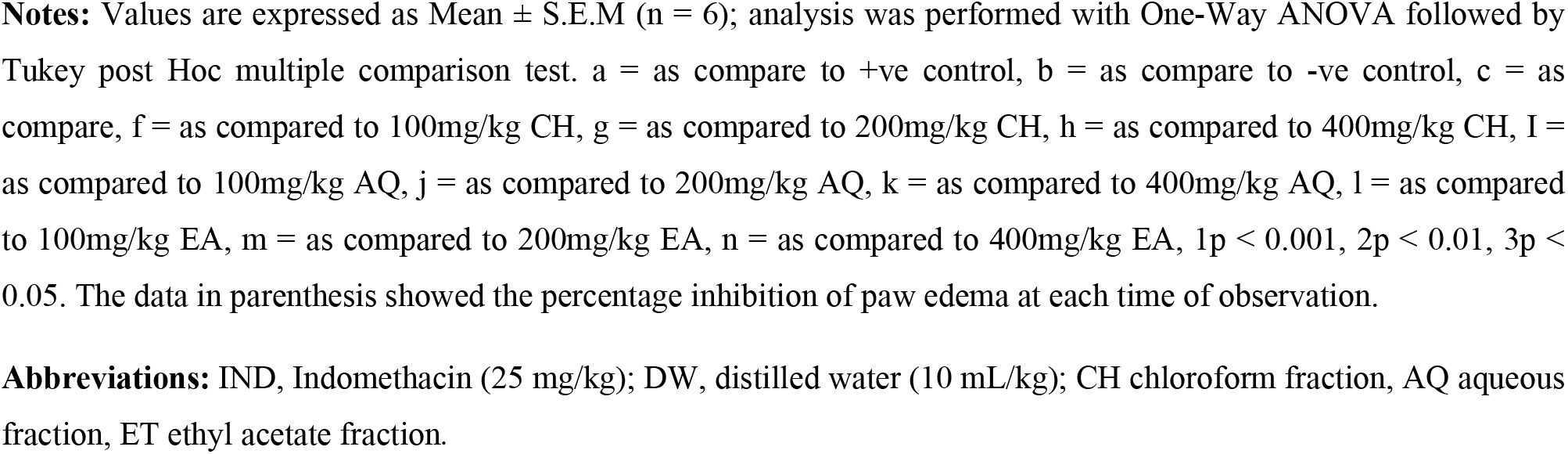
Effect of solvent fractions of EK on Carrageenan-Induced Paw Edema Model in Mice

| Change in Edema Volume (mL) Mean ± SEM | | | | | |
| --- | --- | --- | --- | --- | --- |
| Groups | 0hr | 1hr | 2hr | 3hr | 4hr |
| IND<br>25mg/kg | 0.56 ±<br>0.03 | 0.3 ± 0.01b<sup>1</sup>c<sup>3</sup>d<sup>2</sup><br>(45.78) | 0.25 ± 0.02b<sup>1</sup>c<sup>2</sup><br>(52.1) | 0.25 ± 0.01b<sup>1</sup><br>(59.23) | 0.2±<br>0.01b<sup>1</sup>c<sup>1</sup>d<sup>2</sup>(63.13) |
| DW<br>10ml/kg | 0.53 ±<br>0.03 | 0.55 ± 0.01 | 0.53 ± 0.01 | 0.62 ± 0.03 | 0.53 ± 0.1 |
| EA 100<br>mg/kg | 0.52 ±<br>0.02 | 0.4 ±<br>0.02a<sup>2</sup>b<sup>1</sup>(28.01) | 0.4 ± 0.03b<sup>1</sup>(31.0) | 0.34 ±<br>0.01b<sup>1</sup>(44.77) | 0.33±<br>0.02a<sup>1</sup>b<sup>3</sup>f<sup>2</sup>h<sup>1</sup>(38.75) |
| EA 200<br>mg/kg | 0.5 ±<br>0.02 | 0.36 ±<br>0.02b<sup>1</sup>(34.33) | 0.32 ±<br>0.02b<sup>1</sup>(39.12) | 0.31 ±<br>0.01b<sup>1</sup>(49.9) | 0.27 ± 0.02b<sup>1</sup>f<sup>2</sup>(50) |
| EA 400<br>mg/kg | 0.53 ±<br>0.01 | 0.38 ±<br>0.02b<sup>1</sup>(31.32) | 0.32 ±<br>0.02b<sup>1</sup>(37.54) | 0.29 ±<br>0.02b<sup>1</sup>(52.28) | 0.25 ± 0.01b<sup>1</sup>f<sup>1</sup>(51.25) |
| AQ 100<br>mg/kg | 0.53 ±<br>0.01 | 0.45 ±<br>0.02a<sup>1</sup>b<sup>1</sup>(18.67) | 0.44 ±<br>0.02a<sup>1</sup>b<sup>2</sup>(16.4) | 0.42 ±<br>0.02a<sup>1</sup>b<sup>1</sup>(32.44) | 0.41 ±<br>0.02a<sup>1</sup>b<sup>1</sup>j<sup>2</sup>k<sup>1</sup>(23.13) |
| AQ 200<br>mg/kg | 0.53 ±<br>0.02 | 0.40 ±<br>0.02a<sup>1</sup>b<sup>1</sup>(27.71) | 0.37 ±<br>0.02a<sup>1</sup>b<sup>1</sup>(29.97) | 0.35 ±<br>0.01b<sup>1</sup>(43.16) | 0.34 ±<br>0.01a<sup>1</sup>b<sup>1</sup>i<sup>3</sup>(36.87) |
| AQ 400<br>mg/kg | 0.54 ±<br>0.02 | 0.41 ±<br>0.01a<sup>1</sup>b<sup>1</sup>(36.75) | 0.35 ±<br>0.01a<sup>2</sup>b<sup>1</sup>i<sup>2</sup>(38.17) | 0.33 ±<br>0.02b<sup>1</sup>(45.31) | 0.29 ±<br>0.02a<sup>1</sup>b<sup>1</sup>i<sup>1</sup>(45.63) |
| CH 100<br>mg/kg | 0.55 ±<br>0.01 | 0.50 ±<br>0.01a<sup>1</sup>(9.63) | 0.46 ±<br>0.01b<sup>1</sup>n<sup>3</sup>(13.56) | 0.44 ±<br>0.02a<sup>1</sup>b<sup>1</sup>(28.69) | 0.42 ±<br>0.01a<sup>1</sup>b<sup>1</sup>n<sup>3</sup>(21.87) |
| CH 200<br>mg/kg | 0.57 ±<br>0.01 | 0.47 ± 0.02a<sup>1</sup>(8.0) | 0.43 ±<br>0.02a<sup>1</sup>b<sup>2</sup>(19.24) | 0.42 ±<br>0.01a<sup>1</sup>b<sup>1</sup>(31.90) | 0.39 ± 0.01a<sup>1</sup>b<sup>1</sup>(26.25) |
| CH 400<br>mg/kg | 0.55 ±<br>0.02 | 0.43 ±<br>0.01a<sup>1</sup>b<sup>1</sup>l<sup>3</sup>(22.89) | 0.38 ±<br>0.01a<sup>1</sup>b<sup>1</sup>l<sup>3</sup>(27.44) | 0.36 ±<br>0.03a<sup>3</sup>b<sup>1</sup>(41.56) | 0.35± 0.01a<sup>1</sup>b<sup>1</sup>l<sup>3</sup>(35.0) |

**Notes:** Values are expressed as Mean ± S.E.M (n = 6); analysis was performed with One-Way ANOVA followed by Tukey post Hoc multiple comparison test. a = as compare to +ve control, b = as compare to -ve control, c = as compare, f = as compared to 100mg/kg CH, g = as compared to 200mg/kg CH, h = as compared to 400mg/kg CH, I = as compared to 100mg/kg AQ, j = as compared to 200mg/kg AQ, k = as compared to 400mg/kg AQ, l = as compared to 100mg/kg EA, m = as compared to 200mg/kg EA, n = as compared to 400mg/kg EA, 1p < 0.001, 2p < 0.01, 3p < 0.05. The data in parenthesis showed the percentage inhibition of paw edema at each time of observation.
**Abbreviations:** IND, Indomethacin (25 mg/kg); DW, distilled water (10 mL/kg); CH chloroform fraction, AQ aqueous fraction, ET ethyl acetate fraction.

### Hot plate Method

The hot plate method was conducted to assess the central analgesic effects of solvent fractions of the roots of *E*.*kebericho* in a mice model. In this model, the standard drug (morphine 10mg/kg) and the different test doses of EA, AQ and CH fractions of the roots of *E. kebericho* (400mg/kg, 200mg/kg, and 100mg/kg) showed significant central analgesics activity (p < 0.001, p < 0.01 and p < 0.05) starting from 30 minutes of observation by delaying reaction time as compared to the negative control group **(Table 2)**. In all cases, the highest doses (400mg/kg) produced statistically significant central analgesic activities (p < 0.001) by delaying the reaction time at the 90^th^time of observation as compared to the negative control group. The EA fraction at all test doses (100mg/kg, 200mg/kg and 400mg/kg) showed a significant (p < .001, p < 0.01 and p < 0.05 respectively) analgesic activity starting from 30minutes of observation onwards as compared with the negative control (**Table 2**). As summarized in **table 3** Among the three test doses of EA fractions, the maximum dose (400mg/kg) produced high analgesic activity at the 90^th^minute of observations (60%) as compared to the minimum and middle doses (100mg/kg and 200mg/kg) which produced (30.67% and 42.96%) analgesic activities respectively. Likewise, the test doses of AQ fraction (100mg/kg, 200mg/kg, and 400mg/kg) produce a high percentage of analgesic activities at 90minut of observation with the respective values of 35.08%, 32.34%, and 41.47% respectively. The test doses (100mg/kg, 200mg/kg, and 400mg/kg) of CH fraction on the other hand produced maximum analgesic activities at 120minut of observation with the respective values of 32.38%, 34.58%, and 43.00% respectively.

**Table 2.**
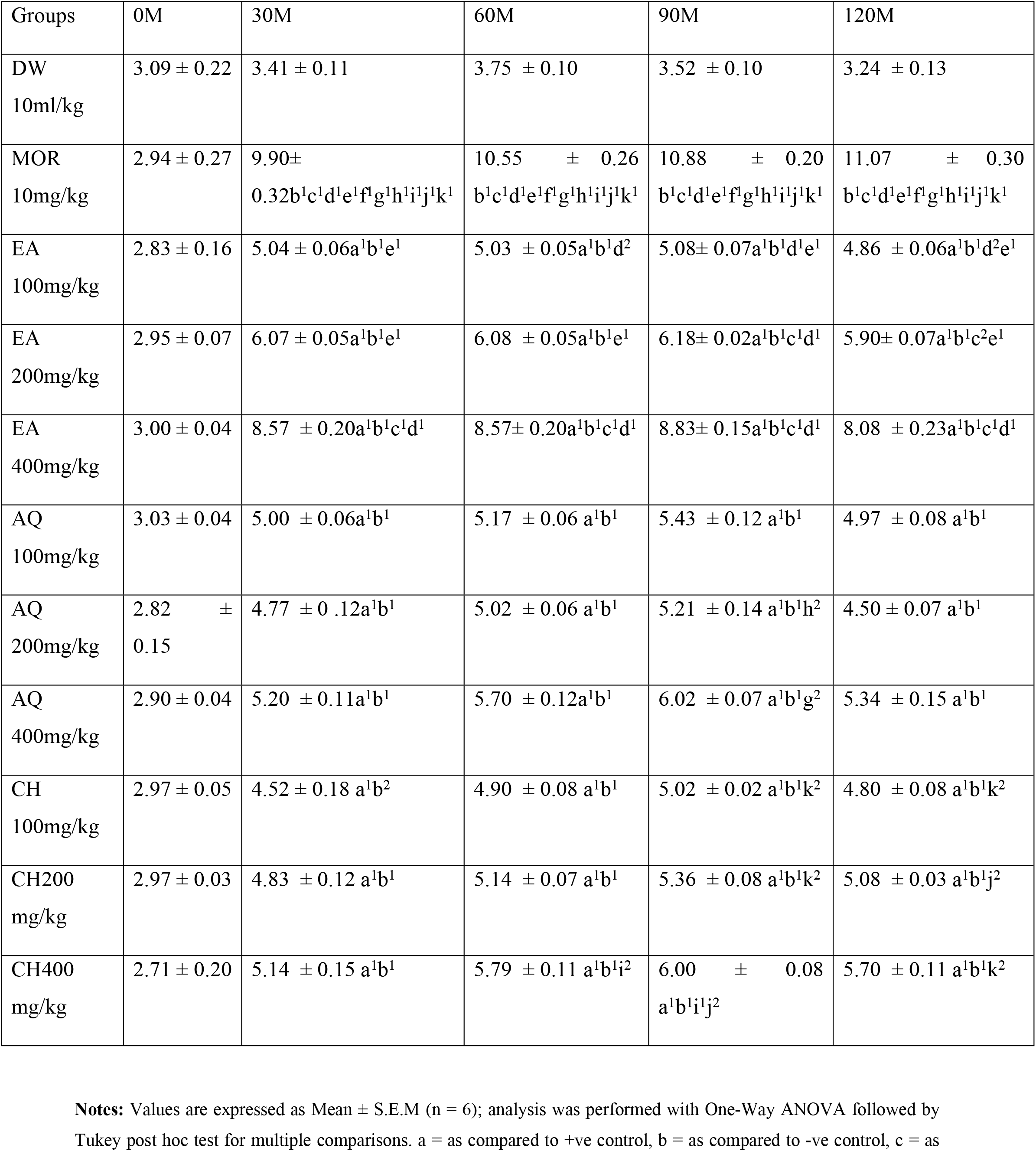
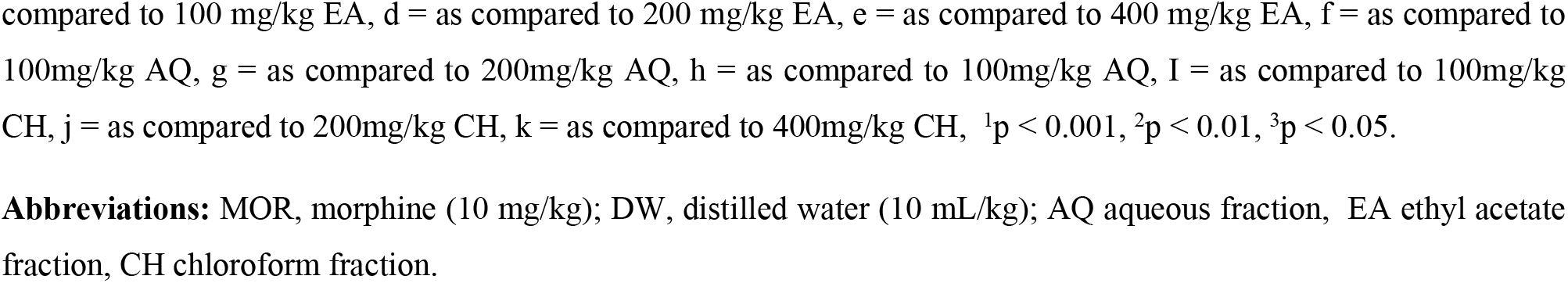
Effects of Solvent Fractions of the Roots of *Echinops kebericho* on Hot Plate Latency Time in Mice

| Groups | 0M | 30M | 60M | 90M | 120M |
| --- | --- | --- | --- | --- | --- |
| DW<br>10ml/kg | 3.09 ± 0.22 | 3.41 ± 0.11 | 3.75 ± 0.10 | 3.52 ± 0.10 | 3.24 ± 0.13 |
| MOR<br>10mg/kg | 2.94 ± 0.27 | 9.90±<br>0.32b<sup>1</sup>c<sup>1</sup>d<sup>1</sup>e<sup>1</sup>f<sup>1</sup>g<sup>1</sup>h<sup>1</sup>i<sup>1</sup>j<sup>1</sup>k<sup>1</sup> | 10.55 ± 0.26<br>b<sup>1</sup>c<sup>1</sup>d<sup>1</sup>e<sup>1</sup>f<sup>1</sup>g<sup>1</sup>h<sup>1</sup>i<sup>1</sup>j<sup>1</sup>k<sup>1</sup> | 10.88 ± 0.20<br>b<sup>1</sup>c<sup>1</sup>d<sup>1</sup>e<sup>1</sup>f<sup>1</sup>g<sup>1</sup>h<sup>1</sup>i<sup>1</sup>j<sup>1</sup>k<sup>1</sup> | 11.07 ± 0.30<br>b<sup>1</sup>c<sup>1</sup>d<sup>1</sup>e<sup>1</sup>f<sup>1</sup>g<sup>1</sup>h<sup>1</sup>i<sup>1</sup>j<sup>1</sup>k<sup>1</sup> |
| EA<br>100mg/kg | 2.83 ± 0.16 | 5.04 ± 0.06a<sup>1</sup>b<sup>1</sup>e<sup>1</sup> | 5.03 ± 0.05a<sup>1</sup>b<sup>1</sup>d<sup>2</sup> | 5.08± 0.07a<sup>1</sup>b<sup>1</sup>d<sup>1</sup>e<sup>1</sup> | 4.86 ± 0.06a<sup>1</sup>b<sup>1</sup>d<sup>2</sup>e<sup>1</sup> |
| EA<br>200mg/kg | 2.95 ± 0.07 | 6.07 ± 0.05a<sup>1</sup>b<sup>1</sup>e<sup>1</sup> | 6.08 ± 0.05a<sup>1</sup>b<sup>1</sup>e<sup>1</sup> | 6.18± 0.02a<sup>1</sup>b<sup>1</sup>c<sup>1</sup>d<sup>1</sup> | 5.90± 0.07a<sup>1</sup>b<sup>1</sup>c<sup>2</sup>e<sup>1</sup> |
| EA<br>400mg/kg | 3.00 ± 0.04 | 8.57 ± 0.20a<sup>1</sup>b<sup>1</sup>c<sup>1</sup>d<sup>1</sup> | 8.57± 0.20a<sup>1</sup>b<sup>1</sup>c<sup>1</sup>d<sup>1</sup> | 8.83± 0.15a<sup>1</sup>b<sup>1</sup>c<sup>1</sup>d<sup>1</sup> | 8.08 ± 0.23a<sup>1</sup>b<sup>1</sup>c<sup>1</sup>d<sup>1</sup> |
| AQ<br>100mg/kg | 3.03 ± 0.04 | 5.00 ± 0.06a<sup>1</sup>b<sup>1</sup> | 5.17 ± 0.06 a<sup>1</sup>b<sup>1</sup> | 5.43 ± 0.12 a<sup>1</sup>b<sup>1</sup> | 4.97 ± 0.08 a<sup>1</sup>b<sup>1</sup> |
| AQ<br>200mg/kg | 2.82 ±<br>0.15 | 4.77 ± 0.12a<sup>1</sup>b<sup>1</sup> | 5.02 ± 0.06 a<sup>1</sup>b<sup>1</sup> | 5.21 ± 0.14 a<sup>1</sup>b<sup>1</sup>h<sup>2</sup> | 4.50 ± 0.07 a<sup>1</sup>b<sup>1</sup> |
| AQ<br>400mg/kg | 2.90 ± 0.04 | 5.20 ± 0.11a<sup>1</sup>b<sup>1</sup> | 5.70 ± 0.12a<sup>1</sup>b<sup>1</sup> | 6.02 ± 0.07 a<sup>1</sup>b<sup>1</sup>g<sup>2</sup> | 5.34 ± 0.15 a<sup>1</sup>b<sup>1</sup> |
| CH<br>100mg/kg | 2.97 ± 0.05 | 4.52 ± 0.18 a<sup>1</sup>b<sup>2</sup> | 4.90 ± 0.08 a<sup>1</sup>b<sup>1</sup> | 5.02 ± 0.02 a<sup>1</sup>b<sup>1</sup>k<sup>2</sup> | 4.80 ± 0.08 a<sup>1</sup>b<sup>1</sup>k<sup>2</sup> |
| CH200<br>mg/kg | 2.97 ± 0.03 | 4.83 ± 0.12 a<sup>1</sup>b<sup>1</sup> | 5.14 ± 0.07 a<sup>1</sup>b<sup>1</sup> | 5.36 ± 0.08 a<sup>1</sup>b<sup>1</sup>k<sup>2</sup> | 5.08 ± 0.03 a<sup>1</sup>b<sup>1</sup>j<sup>2</sup> |
| CH400<br>mg/kg | 2.71 ± 0.20 | 5.14 ± 0.15 a<sup>1</sup>b<sup>1</sup> | 5.79 ± 0.11 a<sup>1</sup>b<sup>1</sup>i<sup>2</sup> | 6.00 ± 0.08<br>a<sup>1</sup>b<sup>1</sup>i<sup>1</sup>j<sup>2</sup> | 5.70 ± 0.11 a<sup>1</sup>b<sup>1</sup>k<sup>2</sup> |

**Notes:** Values are expressed as Mean ± S.E.M (n = 6); analysis was performed with One-Way ANOVA followed by Tukey post hoc test for multiple comparisons. a = as compared to +ve control, b = as compared to -ve control, c = as
compared to 100 mg/kg EA, d = as compared to 200 mg/kg EA, e = as compared to 400 mg/kg EA, f = as compared to 100mg/kg AQ, g = as compared to 200mg/kg AQ, h = as compared to 100mg/kg AQ, I = as compared to 100mg/kg CH, j = as compared to 200mg/kg CH, k = as compared to 400mg/kg CH, <sup>1</sup>p < 0.001, <sup>2</sup>p < 0.01, <sup>3</sup>p < 0.05.
**Abbreviations:** MOR, morphine (10 mg/kg); DW, distilled water (10 mL/kg); AQ aqueous fraction, EA ethyl acetate fraction, CH chloroform fraction.

**Table 3.** Percentage analgesic activities of solvent fractions of the roots of *Echinops kebericho* in thermal-induced latency in mice

| Percentage analgesic activity |  |  |  |  |
| --- | --- | --- | --- | --- |
| Groups | 30M | 60M | 90M | 120M |
| DW 10ml/kg | ----- | ----- | ----- | ----- |
| MOR 10mg/kg | 67.68 | 63.41 | 67.61 | 70.75 |
| EA 100mg/kg | 32.18 | 25.55 | 30.68 | 33.33 |
| EA 200mg/kg | 40.79 | 38.31 | 42.96 | 45.08 |
| EA 400mg/kg | 27.62 | 56.27 | 60.11 | 59.90 |
| AQ 100mg/kg | 31.56 | 27.50 | 35.08 | 34.83 |
| AQ 200mg/kg | 28.41 | 25.33 | 32.34 | 27.92 |
| AQ 400mg/kg | 34.18 | 34.08 | 41.48 | 39.36 |
| CH 100mg/kg | 24.32 | 23.45 | 29.76 | 32.38 |
| CH200 mg/kg | 29.23 | 27.12 | 34.23 | 35.48 |
| CH400 mg/kg | 40.25 | 35.24 | 41.20 | 42.96 |

Analysis was performed with One-Way ANOVA followed by Tukey post hoc multiple comparison test. Data were expressed in mean ± SEM. N = 6.
**Abbreviations:** MOR, morphine (10 mg/kg); DW distilled water (10 mL/kg); EA ethyl Acetate fraction AQ Aqueous fraction; CH Chloroform fraction.

### Anti-inflammatory activity

#### Carrageenan-Induced Paw Edema

This model was conducted to assess the anti-inflammatory properties of solvent fractions of the roots of *E .kebericho* in the acute phase of inflammation. A sub plantar injection of 0.05 mL of 1% carrageenan into the mice hind paw resulted in a gradual increase in paw thickness that peaked after 3 hours of induction in the negative control group. All test doses of ET fraction (100mg/kg, 200mg/kg, and 400mg/kg) of the roots of *E*.*kebericho* exhibited statistically significant inhibition of paw thickness starting from 1hr of observation (p< 0.05, p < 0.01, and 0.001 respectively) and the effects persisted till the fourth hour of observation (p < 0.01 and p < 0.001) post carrageenan induction as compared to the negative control group. On the other hand, AQ and CH fractions exhibited statistical significant (p < 0.05) paw edema inhibition starting from 1 hr of observation onwards at the doses of 400mg/kg, whereas 100mg/kg and 200mg/kg doses of AQ and CH fractions did not show statistical significant (p > 0.05) anti-inflammatory activities.

The maximum percent of mice paw edema inhibition of the standard drug, the extract, and the solvent fractions of *E*.*kebericho* was observed at the 4^th^time of observation with the respective percentage values of 63.15% for the standard drug, 47.5% for *AS* 100mg/kg, 50% for *AS* 56.87% for *AS* 400mg/kg. Likewise, CH fraction exhibited a high percentage of anti-inflammatory activities at the 4^th^ time of observation in dose-dependent manner with the respective values of 44.77%, 50.00%, and 52.3% for 100mg/kg, 200mg/kg, and 400mg/kg respectively (table 1). On the other hand, AQ fraction showed a high percentage of edema inhibition at the doses of 100mg/kg and 200mg/kg at the 3^rd^time of observation with respective values of 32.44% and 43.16% whereas the highest dose (400mg/kg) of AQ fraction showed maximum edema inhibition at the 4^th^time of observation (45.63%). The NB fraction showed the maximum percentage of edema inhibition in all test doses employed (100mg/kg, 200mg/kg, and 400mg/kg) at the 3^rd^ time of observation with values of 28.70%, 31.90%, and 41.56% respectively (table1)

### Cotton Wool Induced Granuloma

This method was performed to detect the anti-inflammatory potentials of solvent fractions of the roots of EK in chronic inflammatory models. Under this method EA fraction at all test doses employed (100mg/kg, 200mg/kg and 400mg/kg) significantly (p < 0.05, p < 0.01, p < 0.001 respectively) prevented the formation of inflammatory exudates and granuloma mass as compared the negative control group. On the other hand AQ fraction at the doses of 200mg/kg and 400mg/kg significantly (p < 0.05 and p < 0.01 respectively) prevented the formation of inflammatory exudates and granuloma mass as compared to the negative control group, whereas CH at the dose of 400mg/kg significantly (p < 0.01) prevented the formation of inflammatory exudates and granuloma mass as compared the negative control group.

## Discussion

Despite the availability of standard drugs for the pharmacological management of pain and inflammation, these issues continue to be a global public health concern that affects over 80% of the population. Furthermore, current anti-pain and anti-inflammatory medications have a variety of untoward effects and toxicities. Ethiopian folk medicine practitioners employ traditional herbal treatments to relieve pain and inflammation (5, 7, 45). Considering the socioeconomic consequences of pain and inflammation, as well as having the knowledge of potential herbal medicines derived from traditionally claimed plants, the need for searching for effective anti-pain and anti-inflammatory drugs with minimal side effects derived from traditional medicinal plants appears reasonable. In this sense, finding medicinal plants that have been widely used in the community to treat various pain problems and inflammation is a critical topic. *Echinops kebericho* M. is among the widely used traditional medicinal plants in Ethiopian folk medicine for treating pain, and different inflammatory conditions (57, 60). It is also reported that 80% of methanol crude extracts of the roots of *E. kebericho* possessed significant analgesic activities on both chemical-induced peripheral and thermal-induced central pain. Moreover, the crude root extract of this plant showed significant anti-inflammatory activities in acute and sub-acute inflammatory models (76). Therefore this study attempted to further evaluate the analgesic and anti-inflammatory activities of the solvent fractions of this plant.

By examining the biological activity of crude extracts or solvent fractions of traditionally claimed herbs, several studies have verified the use of analgesic and anti-inflammatory medicinal plants. The crude extract of the study plant’s roots was tested for analgesic and anti-inflammatory properties, while the current study looked at the anti-pain and anti-inflammation effects of solvent fractions of the study (3, 4, 7). The acetic acid-induced writhing method was used to evaluate the peripheral analgesic effects of the solvent fractions of the roots of *E. kebericho*. Increased levels of prostaglandins (PGE2 and PGF2) in peritoneal fluids, as well as lipooxygenase (LOX) products, have all been linked to the constriction response of the abdomen, which entails the release of arachidonic acid (AA) from tissue phospholipids via cyclooxygenase (COX (76). The peripheral chemo-sensitive nociceptors, which are largely responsible for producing inflammatory pain, are activated and sensitized by PGs, causing abdominal constriction.

The hot plate method, on the other hand, was employed for the assessment of the central analgesic activities of solvent fractions of the roots of *E. kebericho*. The effectiveness of agents in this model shows analgesic activity by a central mechanism acting through opioid receptors and is highly correlated with relief of human pain perception (60).

The acetic acid-induced writhing test, also commonly known as the abdominal contraction test, is used for a reliable and rapid evaluation of the peripheral analgesic action of natural products. The test has long been used as a screening tool to evaluate the antinociceptive and anti-inflammatory properties of new substances. Pain sensation in this writhing method is elicited by acetic acid is believed to act indirectly by inducing the release of prostaglandins as well as lipooxygenase products into the peritoneum which stimulate the nociceptive neurons on the sensory nerve fibers. The acetic acid-induced writhing test is a model of visceral pain (1, 7, 8). It is very sensitive and able to detect antinociceptive effects of compounds at dose levels that may appear inactive in other methods. In this method, EA fraction at the middle and maximum doses (200mg/kg and 400mg/kg) significantly (p < 0.01 and p < 0.001 respectively) showed peripheral analgesic activity by reducing the number of writhing with the respective percentage values of 26.86% and 33.08% as compared with the negative control group whereas the lowest dose (100mg/kg) of EA fraction did not show significant (p > 0.05) analgesic activity as compared with the negative control group. This finding confirmed the fact that the active phytoconstituents that have peripheral antinociceptive activities concentrated with 200mg/kg and 400mg/kg doses of EA fraction. On the other hand, AQ fraction at all test doses employed (100mg/kg/ 200mg/kg and 400mg/kg) produced significant (p < 0.05, p < 0.01, and p < 0.001 respectively) peripheral analgesic effects by reducing the numbers of writhing from 15 minutes of observation onwards in a dose dependant manner with the respective percentage values of 9.5%, 22.95% and 29.35% as compared with the negative control group. This finding assured that most of the active metabolites that have activities for peripheral analgesic activities are better dissolved with distilled water and present on all test doses employed in various concentrations. The CH fraction, on the other hand, produced a statistically significant (p < 0.001) peripheral analgesic effect at the dose of 400mg/kg, whereas 100mg/kg and 200mg/kg doses of CH fraction did not produce a statistically significant (p > 0.05) peripheral analgesic activities as compared with the negative control group. This finding revealed that the analgesic active phytoconstituents are present at a concentration such that they can inhibit different pain mediators at 400mg/kg dose of CH fraction.

Inhibiting the synthesis and release of different endogenous inflammatory mediators, as well as suppressing the sensitivity of peripheral nociceptors in peritoneal free nerve endings to chemical-induced pain, could be the mechanism by which each solvent fraction elicited peripheral antinociceptive effects in this model. These hypothesized processes are in keeping with the concept that any medication that reduces the amount of whriting will produce analgesia by reducing the creation and release of PGs, as well as inhibiting the transmission of peripheral pain (76).

The second model used was the hot plate method, which is usually used to detect the central analgesic effects of natural products. The detection of opiate (narcotic) analgesics is well-established using this model, which depends on nociceptive responses to temperature stimuli. Analgesics that act on the central nervous system have this feature. Therefore, the result obtained under this model revealed that the solvent fractions of the roots of *E. kebericho* have central analgesic activities which resemble centrally acting opiates. Because of its susceptibility to strong analgesics and little tissue damage, this model was used to assess the extract’s central analgesic potential. A cutoff period of 15 seconds was used to limit the length of time the mouse spent on the hot plate. The model also saves time and ensures that measurements are precise.

Under this model, *E. kebericho* solvent fractions showed a significant (p < 0.05, p < 0.01, and 0.001) central analgesic by increasing the reaction time starting from 30 min of observation onwards as compared to the negative control. As summarized on table 3, the maximum percentage of analgesic activity was observed with high doses of EA fraction (60.11%) at 90 minutes of observation followed by, 59.9% at 120 minutes of observation, 56.28% at 60 minutes of observation, and 27.62% at 30 minutes of observation as compared to the standard drug morphine. The solvent fraction’s central analgesic effects may be mediated by stimulating the periaqueductal gray matter (PAG), which releases endogenous peptides (i.e., endorphin or enkephalin). These endogenous peptides descend the spinal cord and act as inhibitors of pain impulse transmission at the dorsal horn synapse or by peripheral processes that inhibit PG, leukotriene, and other endogenous chemicals implicated in central pain transmission (4). Alkaloids, flavonoids, saponin, tannins phenolic substances, glycosides, coumarins, and triterpenoids chemical elements are found in plants that have analgesic and anti-inflammatory effects. Tannins, flavonoids, and saponins are well known for their capacity to reduce pain perception and have anti-inflammatory activities by inhibiting enzymes implicated in inflammation, particularly those involved in the arachidonic acid metabolic pathway and prostaglandin formation. Tannins may influence the inflammatory response by scavenging free radicals and inhibiting iNOS in macrophages. Saponins, on the other hand, work by inhibiting NO, which reduces pain and inflammation. Alkaloids, polyphenols, saponins, phytosterols, carotenoids, lignans, sesquiterpene alcohols, acetylenic and thiophene chemicals, terpenoids, and essential oil have all been found in previous phytochemical screenings on *EK*. Preliminary phytochemical screening revealed that the root extract of *EK* included secondary metabolites such as saponin, tannin, alkaloids, phenols, flavonoids, glycosides, and steroids. As a result, the presence of these aforementioned and currently recognized phytoconstituents may be responsible for the extract’s analgesic properties (15, 16, 21, 60). So the central analgesic activities of solvent fractions of the roots of EK are assumed due to the presence of the aforementioned active phytoconstituents.

The effects of the solvent fractions on chronic inflammation were tested using the cotton pellet-induced granuloma model. This model is frequently used to evaluate chronic inflammation’s transudative and proliferative components. The amount of exudative material correlates with the weight of wet cotton pellets, while the amount of granulomatous tissue correlates with the weight of dry pellets. According to the findings, EA fraction at all test doses employed, AQ fraction at the middle dose, and CH fraction at the highest dose efficiently prevented the formation of granulomas, implying that each they have anti-inflammatory properties.

Alkaloids, flavonoids, saponin, tannins phenolic substances, glycosides, coumarins, and triterpenoids chemical elements are found in plants that have analgesic and anti-inflammatory effects (45, 57). Tannins, flavonoids, and saponins are well known for their capacity to reduce pain perception and have anti-inflammatory activities by inhibiting enzymes implicated in inflammation, particularly those involved in the arachidonic acid metabolic pathway and prostaglandin formation (3, 4). Tannins may influence the inflammatory response by scavenging free radicals and inhibiting iNOS in macrophages. Saponins, on the other hand, work by inhibiting NO, which reduces pain and inflammation (34, 34). The presence of active phytoconstituents from each solvent fraction contributes to the anti-inflammatory properties of the plant.

## Conclusions and Recommendations

In conclusion, the solvent fractions of the study plant exhibited peripheral analgesic activities and central pain inhibition potentials. The studied plant also showed good anti-inflammatory activity in the chronic inflammatory model. Because of the presence of active secondary metabolites such as alkaloids, flavonoids, saponins, terpenoids, tannins, and essential oils, which have been repeatedly reported to have analgesic and anti-inflammatory properties, these findings could imply the solvent fractions were involved in inhibiting various endogenous inflammatory mediators, pain transmission, and mediators. The research findings provide scientific proof for the traditional claimed usage of *Echinops kebericho* M. in Ethiopian folk medicine for painful illnesses and inflammation.

Further investigation should be done on constituent isolation, binding studies, and electrophysiological methods to fully elucidate *E. Kebericho’s* anti-pain and anti-inflammatory mechanisms, as well as specific mechanisms associated with these effects.

## Abbreviations

AF: Aqueous fraction
ASA: Acetyl salicylic acid
CH: chloroform fraction
DW: distilled water
EA: Ethyl acetate fraction
*EK*: *Echinops kebericho Mesfin*
IFN: Interferon
IL: Interleukin
iNOS: inducible nitric oxide synthase
NO: nitric oxide
PGs: Prostaglandin

## Data Availability

All data that are analyzed are available on the hand of the corresponding author upon reasonable request

## Ethical Approval

Ethical clearance was obtained from the Research and Ethics Committee, College of Health Science, Debre Tabor University to conduct the study in an animal model with the reference number CHS 07-105-21.

## Authors’ Contributions

All authors made a significant contribution to the work reported; participated in the conception, study design, execution, and acquisition of data, analysis, and interpretation; took part in drafting, revising, or critically reviewing the article; gave final approval of the version to be published; agreed on the journal to which the article has been submitted; and agreed to be accountable for all aspects of the work. TYT conceived the idea and drafted the proposal. SBD, TGY, GTA, and ZDK prepared and critically reviewed the final manuscript for publication. All authors read and approved the final version of the manuscript.

## Acknowledgments

The authors would like to acknowledge Debre Tabor University.

